# Moderate overexpression of *PROTON GRADIENT REGULATION 5* improves photosynthetic performance and plant growth under fluctuating low light in *Arabidopsis thaliana*

**DOI:** 10.64898/2026.04.21.719829

**Authors:** Keiichiro Tanigawa, Hiromasa Kodama, Yuki Okegawa, Toshiharu Shikanai, Wataru Yamori

## Abstract

Cyclic electron transport (CET) around photosystem I (PSI) is essential for maintaining photosynthetic efficiency by balancing ATP/NADPH production and protecting PSI from photoinhibition. Although the PROTON GRADIENT REGULATION 5 (PGR5)–dependent CET pathway is known to be critical under high or fluctuating light conditions, its role under fluctuating low light remains poorly understood. In natural environments, plants frequently experience prolonged low irradiance interspersed with brief sunflecks, making fluctuating low light a physiologically relevant condition. Here, we investigated *Arabidopsis thaliana* lines with graded *PGR5* expression levels to evaluate the dose-dependent contribution of PGR5 to CET activity, photosynthetic regulation, and growth performance under both low light and fluctuating low light conditions. Moderate increase in the PGR5 protein level enhanced CET activity, accelerated photosynthetic induction, improved PSI protection and increased biomass accumulation under fluctuating low light. In contrast, excessive PGR5 accumulation impaired photosynthetic performance and reduced plant growth, indicating that optimal CET capacity requires precise tuning of PGR5 abundance. These results reveal a non-linear relationship between PGR5 protein levels and photosynthetic performance and demonstrate that moderate enhancement of CET improves plant productivity under fluctuating low light. Our findings highlight the importance of optimizing CET capacity to match dynamic light environments and suggest that fine-tuning PGR5 expression could be a promising strategy for improving crop performance under natural canopy conditions.

**Significance statement:** Moderate increase in the PGR5 improves plant productivity, whereas excessive PGR5 accumulation impaired photosynthetic performance and reduced plant growth. Therefore, optimizing CET capacity by the fine-tuning PGR5 expression is important for improving crop productivity.

## Introduction

Photosynthesis is the fundamental process by which light energy is converted into chemical energy, sustaining almost all life on Earth. In chloroplasts, light energy drives the production of ATP and NADPH, which serve as essential energy sources for the carbon-fixing reactions of the Calvin-Benson cycle (Yamori and Shikanai, 2016). The primary electron transport pathway, known as linear electron transport (LET), involves the transfer of electrons from water via photosystem II (PSII), plastoquinone (PQ), the cytochrome *b*_6_*f* complex (Cyt *b*_6_*f*), plastocyanin (PC), and photosystem I (PSI), ultimately reducing NADP^+^ to NADPH. Coupled with LET, protons are translocated from the stroma into the thylakoid lumen, generating a proton motive force (*pmf*) across the thylakoid membrane that drives ATP synthesis (Allen, 2002).

Despite the central role of LET, it does not satisfy the stoichiometric requirements of the Calvin-Benson cycle, which demands an ATP/NADPH ratio of approximately 1.5 (Allen, 2003). LET alone typically yields a ratio of about 1.29 due to the fixed stoichiometry of proton translocation and ATP synthesis (Hahn et al., 2018). To compensate for this imbalance and to regulate the redox poise of the electron transport chain, plants employ an alternative pathway known as cyclic electron transport (CET) around PSI (Yamori & Shikanai, 2016). CET functions to generate additional *pmf* and ATP without the concomitant production of NADPH, thereby restoring the ATP/NADPH balance and providing photoprotective benefits under fluctuating environmental conditions (Shikanai, 2007).

In angiosperms, CET is mediated through two partially redundant pathways: one dependent on the PROTON GRADIENT REGULATION 5 (PGR5) and the other involving the chloroplast NADH dehydrogenase-like (NDH) complex (Endo et al., 1998; Munekage et al., 2002; DalCorso et al., 2008). Among these, the PGR5-dependent pathway is considered the primary route in most C_3_ plants under normal physiological conditions. Mutants deficient in PGR5 (*pgr5*) exhibit severe photosynthetic defects, including impaired *pmf* formation, reduced non-photochemical quenching (NPQ), and increased photoinhibition of PSI, particularly under fluctuating or high light conditions (Munekage et al., 2002; Suorsa et al., 2012; Yamori et al., 2016; Kobayashi et al., 2024). These phenotypes underline the critical role of PGR5 in maintaining redox balance and protecting the photosynthetic machinery under variable light environments (Tikkanen et al., 2014; Yamori and Shikanai, 2016).

The molecular function of PGR5, however, remains a subject of ongoing investigation. Previously, the most widely accepted hypothesis was that PGR5 does not itself function as an electron carrier but rather plays a regulatory role in controlling the flow of electrons from PSI to the PQ pool, possibly via interaction with PGRL1. The PGR5-PGRL1 complex has been proposed to serve as a ferredoxin-plastoquinone reductase (FQR), catalyzing the reduction of PQ using electrons from ferredoxin (DalCorso et al., 2008). However, recent studies are leading to a re-evaluation of this long-held hypothesis. Rühle et al. (2021) suggested that PGR5 likely functions independently. The role of PGRL1 is related to maintaining the stability of PGR5, specifically by potentially preventing its degradation by another protein, PGRL2. Recent evidence also indicates that the activity of this complex may be tightly regulated by the redox state of the stroma and post-translational modifications, such as thiol modulation (Okegawa et al., 2020).

The physiological importance of CET is particularly pronounced under fluctuating light (FL) conditions, which are prevalent in natural environments due to sunflecks, cloud cover, and leaf movements (Pearcy, 1990; Yamori 2016). Under such conditions, PSI is especially vulnerable to photoinhibition caused by over-reduction of its acceptor side. PGR5-dependent CET has been shown to prevent PSI over-reduction by sustaining an adequate *pmf*, which in turn slows down electron flow into PSI through downregulation of the Cyt *b*_6_*f* complex (Suorsa et al., 2012). This mechanism, known as photosynthetic control, helps maintain PSI activity and protects against photodamage (Kono et al., 2025). In *pgr5* mutants, the loss of CET capacity leads to rapid over-reduction of PSI and subsequent photoinhibition during transitions from low to high light, thereby compromising overall photosynthetic performance and growth (Tikkanen et al., 2014; Yamori et al., 2016).

In addition to its photoprotective function, PGR5-mediated CET is implicated in optimizing photosynthetic induction, a process in which the photosynthetic apparatus gradually activates upon transition from dark or low light to high light. Efficient CET facilitates the establishment of *pmf* and rapid generation of ATP, which are required for activating carbon fixation enzymes and initiating photoprotective responses such as NPQ (Suorsa et al., 2012). Delayed or impaired CET, as observed in *pgr5* mutants, often results in sluggish induction of photosynthesis (Okegawa et al., 2020), reduced energy utilization efficiency, and compromised carbon gain during short light transients.

While the role of PGR5 under high and fluctuating light has been extensively studied, its function under low light conditions remains less understood. Given that natural light environments often include prolonged periods of low irradiance interspersed with brief sunflecks, understanding how PGR5 contribute to photosynthetic performance and plant growth in low light is crucial. Moreover, most studies to date have focused on either loss-of-function mutants or lines with extreme overexpression of *PGR5*, which can result in pleiotropic effects such as altered levels of other thylakoid proteins and aberrant growth phenotypes (Okegawa et al., 2007; Long et al., 2008; Munekage et al., 2008). These limitations hinder a nuanced understanding of how PGR5 protein levels quantitatively affect CET activity and downstream physiological responses.

To address these gaps, it is important to investigate a range of transgenic lines with graded *PGR5* expression levels, allowing the dissection of PGR5’s dose-dependent effects on CET, photosynthetic induction, and plant fitness under varying light environments. Such studies will not only clarify the functional threshold and optimal range of PGR5 activity but also offer insights into how CET contributes to light acclimation strategies in plants. Ultimately, these findings could inform approaches to enhance photosynthetic efficiency and stress resilience in crops by fine-tuning CET capacity through targeted genetic or biochemical interventions.

## Results

### PGR5 protein abundance affects photosynthetic capacity under steady-state conditions

To examine the physiological consequences of altered PGR5 levels, we first quantified the abundance of PGR5 protein in all lines (Figs. 1B, 1C). The PGR5 mutant (*pgr5-5*) accumulated very little PGR5 protein. In contrast, the *PGR5ox 65* line accumulated approximately twice the PGR5 level of wild type (WT), whereas *PGR5ox 5-10* and *PGR5ox 25-10* accumulated approximately three times the WT level. *PGR5ox 20* accumulated half the PGR5 protein level relative to WT, probably due to co-suppression (Fig. 1B, 1C).

**Fig. 1.**
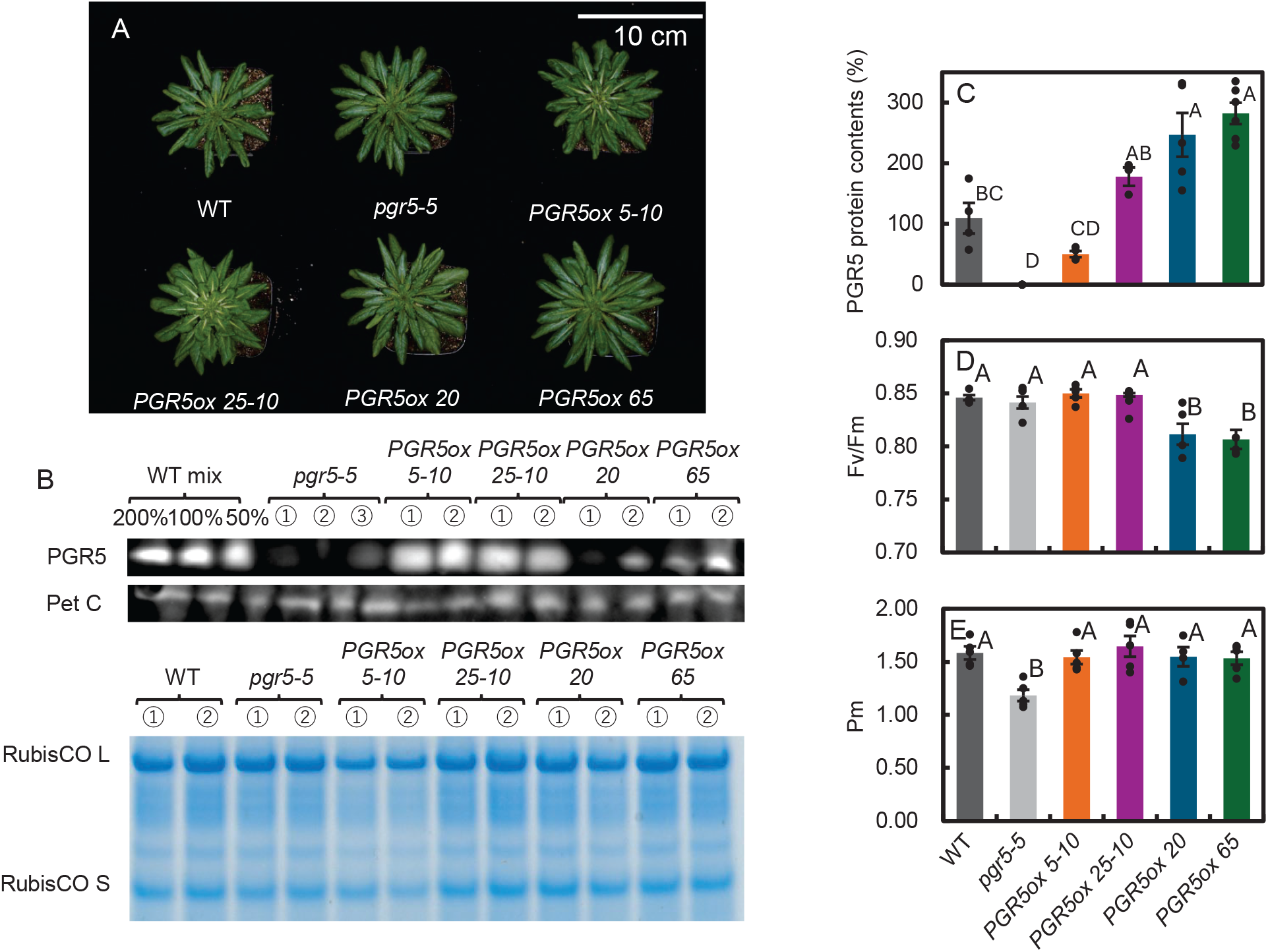
Growth, protein accumulation, steady-state chlorophyll fluorescence and P700 redox state in WT, *pgr5-5, PGR5ox 20, PGR5ox 65, PGR5ox 5-10* and *PGR5ox 25-10*. **A**: Plants were grown in a growth chamber at 120 μmol photons m^−2^ s^−1^ under short-day conditions (10 h of light/14 h of dark) for 6 weeks. **B**: Immunoblot analysis of PGR5 and PetC proteins and SDS-PAGE analysis of RubisCO complex proteins in the leaf. **C**: Relative amount of PGR5 protein upon WT being set at 100%. Data are means ± SE (n□≥□3), and dots indicate individual data points. For each point, different letters indicate a significant difference, as determined by one-way ANOVA with Tukey’s HSD tests (P□<□0.05). **D**: F_v_/F_m_ under the steady state. Data are means ± SE (n□≥□5), and dots indicate individual data points. For each point, different letters indicate a significant difference, as determined by one-way ANOVA with Tukey’s HSD tests (P□<□0.05). **E**: The maximum P_700_ under the steady state. Data are means ± SE (n□≥□5), and dots indicate individual data points. For each point, different letters indicate a significant difference, as determined by one-way ANOVA with Tukey’s HSD tests (P□<□0.05).

In contrast to the large variation in PGR5 abundance, the amounts of the Cyt *b*_6_*f* subunit PetC and RubisCO were similar among all lines (Figs. 1B, S1A, S1B), indicating that alterations in PGR5 levels did not affect the accumulation of these photosynthetic proteins.

Under steady-state conditions, clear genotype-dependent differences in photosystem performance were observed. The two highest PGR5-expressing lines (*PGR5ox 5-10* and *PGR5ox 25-10*) showed significantly lower values of Fv/Fm than the other lines (Fig. 1D), suggesting increased Fo’ due to PQ reduction during dark period (Okegawa et al., 2007), increased photoinhibition or impaired PSII stability at very high PGR5 levels.

In contrast, the maximum oxidizable P700 signal (Pm), which reflects PSI capacity, was significantly lower in the *pgr5-5* mutants than in the other lines (Fig. 1E). These results indicate that both insufficient and excessive accumulation of PGR5 can affect photosystem performance.

### Photosynthetic parameters show distinct relationships with PGR5 abundance under low light

To investigate how PGR5 abundance influences photosynthetic regulation during photosynthetic induction, we analyzed multiple electron transport parameters under low-light (LL) conditions (Fig. 2-1A). The parameters measured included Y(II), Y(I), NPQ, 1−qL, Y(ND), Y(NA), ETRII, ETRI, and the ETRI/ETRII ratio (Figs. 2-1B–2-1L; Figs. S2A–S2G).

**Fig. 2-1.**
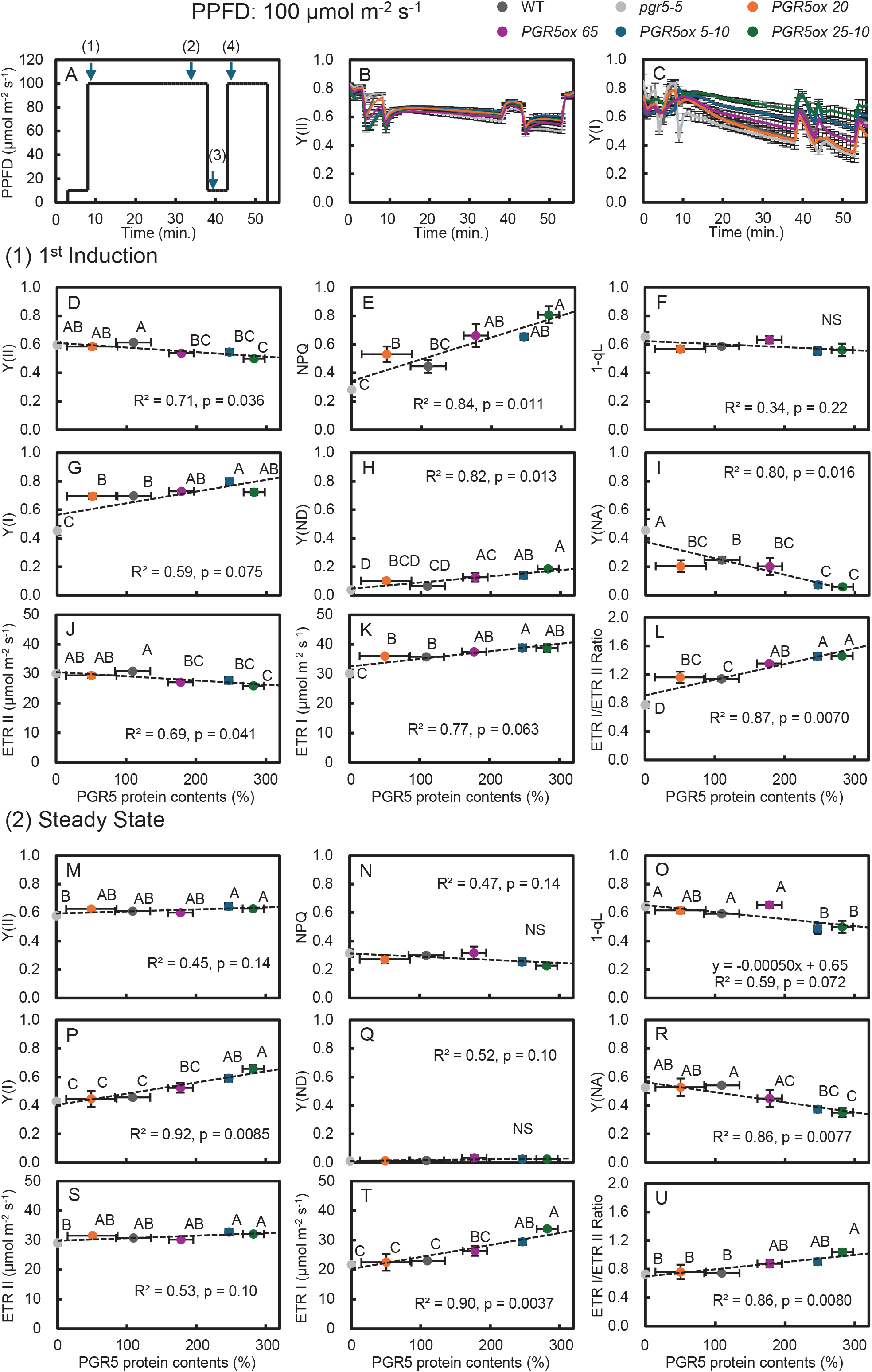
Chlorophyll fluorescence and P700 analysis of a photosynthetic induction response in WT, *pgr5-5, PGR5ox 20, PGR5ox 65, PGR5ox 5-10* and *PGR5ox 25-10* under the LL condition. Time series changes in (**A**) light intensity, (**B**) Y(II), and (**C**) Y(I). A plot of (**D**) Y(II), (**E**) NPQ, (**F**) 1−qL, (**G**) Y(I), (**H**) Y(ND), (**I**) Y(NA), (**J**) ETRII, (**K**) ETRI, and (**L**) ETRI/ETRII ratio as a function of PGR5 protein contents in the 1^st^ induction. A plot of (**M**) Y(II), (**N**) NPQ, (**O**) 1−qL, (**P**) Y(I), (**Q**) Y(ND), (**R**) Y(NA), (**S**) ETRII, (**T**) ETRI, and (**U**) ETRI/ETRII ratio as a function of PGR5 protein contents in the Steady State. Data are means ± SE (n□≥□4). For each point, different letters indicate a significant difference, as determined by one-way ANOVA with Tukey’s HSD tests (P□<□0.05). The regression line was obtained using the method of least squares and evaluated by calculating the R^2^ and p-value based on Pearson’s correlation coefficient.

Overall time-course responses revealed line-dependent patterns of electron transport regulation. In particular, the *PGR5ox 65* line frequently displayed distinct responses compared with the highest-expressing lines (*PGR5ox 5-10* and *PGR5ox 25-10*), especially in NPQ, 1−qL, and Y(NA) (Figs. S2A–S2D), suggesting that moderate PGR5 accumulation influences photosynthetic reaction differently from excessive PGR5 accumulation.

To further examine these relationships, we analyzed four representative time points during the fluctuating light cycle: the first high-light induction phase (1^st^ induction), the steady-state phase during high-light, the phase immediately after light reduction (resolved), and the second induction phase (2^nd^ induction) (Fig. 2-1A).

During the 1^st^ induction phase, several parameters showed clear relationships with PGR5 abundance. Y(II), Y(NA), and ETRII showed significant inverse correlations with PGR5 levels (Figs. 2-1D, 2-1I, 2-1J), whereas NPQ, Y(I), Y(ND), ETRI, and the ETRI/ETRII ratio showed significant positive correlations (Figs. 2-1E, 2-1G, 2-1H, 2-1K, 2-1L). In most parameters, the largest differences were observed between the pgr5 mutants and the highest-expressing line *PGR5ox 25-10*.

During the steady-state phase, Y(II), Y(I), ETRII, ETRI, and the ETRI/ETRII ratio showed strong positive correlations with PGR5 abundance (Figs. 2-1M, 2-1P, 2-1S–2-1U). Conversely, 1−qL and Y(NA) were negatively correlated with PGR5 levels (Figs. 2-1O, 2-1R).

Similar correlations were observed after the light intensity decreased (resolved phase) and during the 2^nd^ induction phase (Figs. 2-2A–2-2R). Together, these results demonstrate that PGR5 abundance strongly influences the regulation of PSI and PSII electron transport under fluctuating low-light conditions.

**Fig. 2-2.**
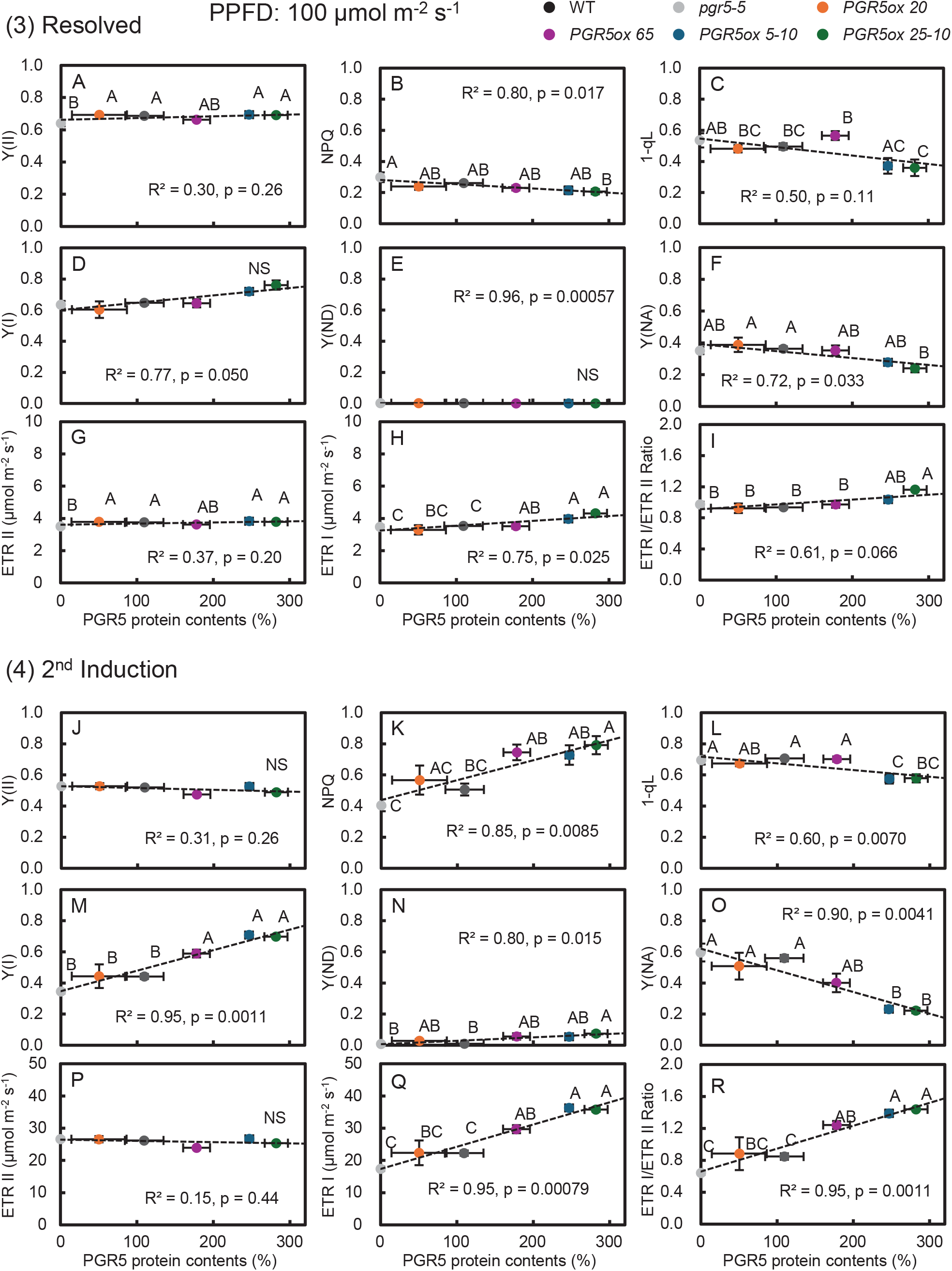
Chlorophyll fluorescence and P700 analysis of a photosynthetic induction response in WT, *pgr5-5, PGR5ox 20, PGR5ox 65, PGR5ox 5-10* and *PGR5ox 25-10* under the LL condition. A plot of (**A**) Y(II), (**B**) NPQ, (**C**) 1−qL, (**D**) Y(I), (**E**) Y(ND), (**F**) Y(NA), (**G**) ETRII, (**H**) ETRI, and (**I**) ETRI/ETRII ratio as a function of PGR5 protein contents in the Resolved. A plot of (**J**) Y(II), (**K**) NPQ, (**L**) 1−qL, (**M**) Y(I), (**N**) Y(ND), (**O**) Y(NA), (**P**) ETRII, (**Q**) ETRI, and (**R**) ETRI/ETRII ratio as a function of PGR5 protein contents in the 2^nd^ induction. Data are means ± SE (n□≥□4). For each point, different letters indicate a significant difference, as determined by one-way ANOVA with Tukey’s HSD tests (P□<□0.05). The regression line was obtained using the method of least squares and evaluated by calculating the R^2^ and p-value based on Pearson’s correlation coefficient.

### Photosynthetic regulation under high light is strongly impaired in pgr5-5 mutants

We next examined photosynthetic responses under high-light (HL) conditions using the same set of parameters (Figs. 3-1B–3-1U; Figs. S3A–S3G).

**Fig. 3-1.**
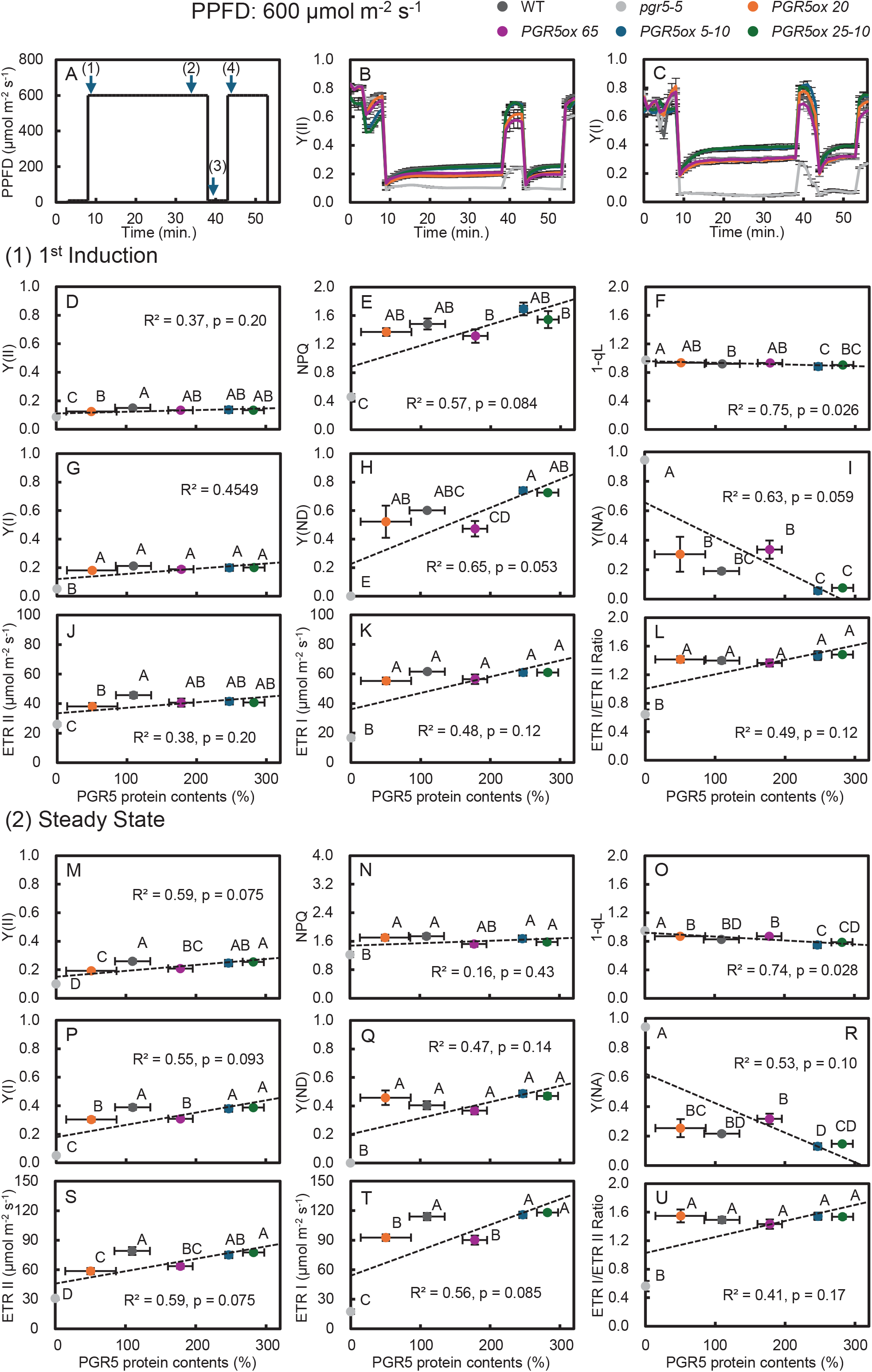
Chlorophyll fluorescence and P700 analysis of a photosynthetic induction response in WT, *pgr5-5, PGR5ox 20, PGR5ox 65, PGR5ox 5-10* and *PGR5ox 25-10* under the HL condition. Time series changes in (**A**) light intensity, (**B**) Y(II), and Y(I). A plot of (**D**) Y(II), (**E**) NPQ, (**F**) 1−qL, (**G**) Y(I), (**H**) Y(ND), (**I**) Y(NA), (**J**) ETRII, (**K**) ETRI, and (**L**) ETRI/ETRII ratio as a function of PGR5 protein contents in the 1^st^ induction. A plot of (**M**) Y(II), (**N**) NPQ, (**O**) 1−qL, (**P**) Y(I), (**Q**) Y(ND), (**R**) Y(NA), (**S**) ETRII, (**T**) ETRI, and (**U**) ETRI/ETRII ratio as a function of PGR5 protein contents in the Steady State. Data are means ± SE (n□≥□4). For each point, different letters indicate a significant difference, as determined by one-way ANOVA with Tukey’s HSD tests (P□<□0.05). The regression line was obtained using the method of least squares and evaluated by calculating the R^2^ and p-value based on Pearson’s correlation coefficient.

During the 1^st^ induction phase, the highest-expressing lines (*PGR5ox 5-10* and *PGR5ox 25-10*) exhibited transient spikes in NPQ, Y(ND) and Y(NA) (Fig. 3-1E, 3-1H, 3-1I, S3A, S3C, S3D).

During steady-state phase, the *pgr5-5* mutants consistently showed the lowest values for Y(II), Y(I), NPQ, ETRII, and ETRI, while exhibiting the highest Y(NA) values (Figs. 3-1B–3-1C; Figs. S3A; S3E–S3F). These results indicate severe limitations in PSI acceptor-side capacity and reduced electron transport in the absence of PGR5.

When the parameters were analyzed at the four representative time points, the *pgr5-5* mutants consistently behaved as outliers. During the 1^st^ induction phase, most parameters—including NPQ, Y(I), Y(ND), ETRII, ETRI, and the ETRI/ETRII ratio—showed significantly lower values in the *pgr5-5* mutants compared with the other lines (Figs. 3-1E–3-1H; Figs. 3-1J–3-1L). In contrast, Y(NA) was significantly higher in the mutants (Fig. 3-1I).

Similar patterns were observed during the steady-state and resolved phases, with the *pgr5-5* mutants consistently showing lower electron transport rates and higher PSI acceptor-side limitation (Figs. 3-1M–3-1U; Figs. 3-2A–3-2I). These results indicate that PGR5 plays a critical role in maintaining electron transport and PSI function under high-light conditions.

**Fig. 3-2.**
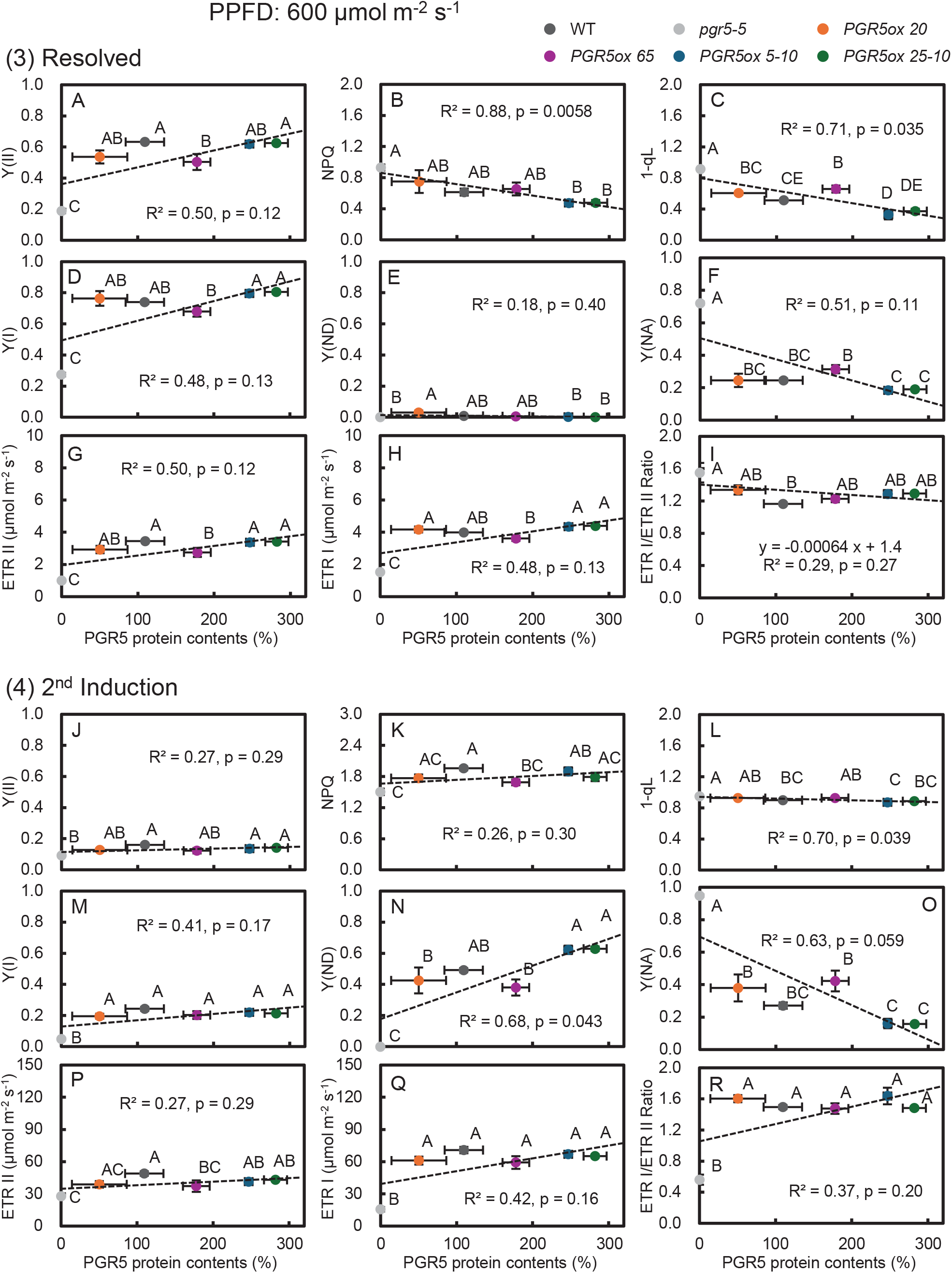
Chlorophyll fluorescence and P700 analysis of a photosynthetic induction response in WT, *pgr5-5, PGR5ox 20, PGR5ox 65, PGR5ox 5-10* and *PGR5ox 25-10* under the HL condition. A plot of (**A**) Y(II), (**B**) NPQ, (**C**) 1−qL, (**D**) Y(I), (**E**) Y(ND), (**F**) Y(NA), (**G**) ETRII, (**H**) ETRI, and (**I**) ETRI/ETRII ratio as a function of PGR5 protein contents in the Resolved. A plot of (**J**) Y(II), (**K**) NPQ, (**L**) 1−qL, (**M**) Y(I), (**N**) Y(ND), (**O**) Y(NA), (**P**) ETRII, (**Q**) ETRI, and (**R**) ETRI/ETRII ratio as a function of PGR5 protein contents in the 2^nd^ induction. Data are means ± SE (n□≥□4). For each point, different letters indicate a significant difference, as determined by one-way ANOVA with Tukey’s HSD tests (P□<□0.05). The regression line was obtained using the method of least squares and evaluated by calculating the R^2^ and p-value based on Pearson’s correlation coefficient.

### Plant growth and photosynthetic traits after growth under different light environments

To determine whether the differences in photosynthetic regulation observed among the lines affected plant growth, plants were cultivated for three weeks under fluctuating or continuous light environments at either LL or HL intensity.

#### 1) Growth responses under low-light conditions

Plant dry weight differed among the lines when plants were grown under fluctuating low light (FLL) (Fig. 4B). In particular, the *PGR5ox 65* line, which accumulated approximately twice the PGR5 protein level of WT, showed significantly higher dry weight than the other lines. In contrast, the remaining lines, including WT, *pgr5-5*, and the two highest-expressing lines (*PGR5ox 5-10* and *PGR5ox 25-10*), showed similar dry weight values under FLL conditions.

**Fig. 4.**
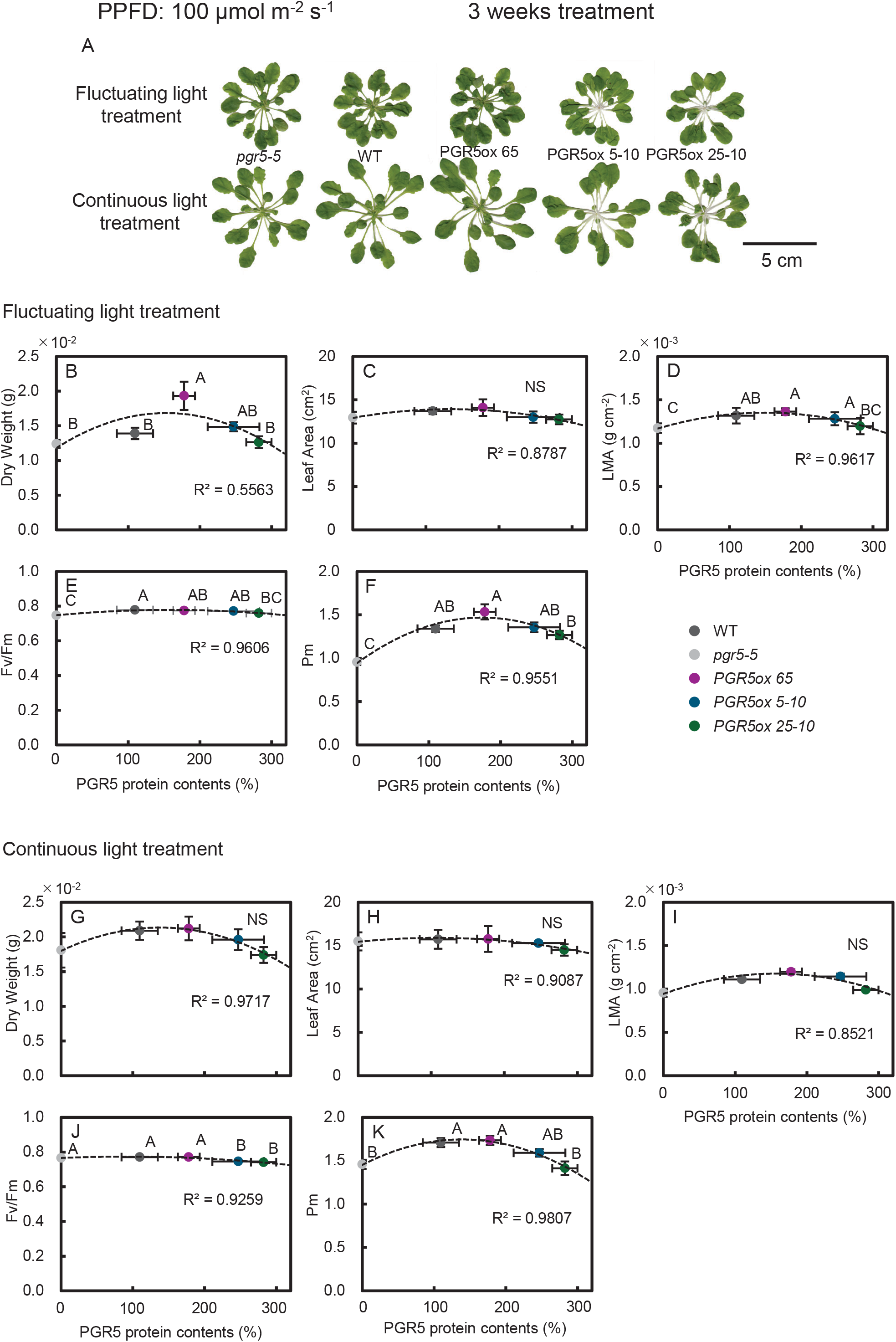
Plants Phenotype and growth analysis under 3 weeks of FLL treatment, chlorophyll fluorescence and P700 redox state under 4 weeks of FLL treatment in WT, *pgr5-5, PGR5ox 65, PGR5ox 5-10* and *PGR5ox 25-10*. **A**: Plants were cut and scanned after 3 weeks FLL treatment. Upper plants were grown under FLL condition and lower plants were grown under CLL condition. A plot of (**B**) Dry Weight, (**C**) Leaf Area, (**D**) LMA and (**E**) Fv/Fm as a function of PGR5 protein contents under FLL condition. A plot of (**F**) Dry Weight, (**G**) Leaf Area, (**H**) LMA and (**I**) Fv/Fm as a function of PGR5 protein contents under CLL condition. Data are means ± SE (n□=□4). For each point, different letters indicate a significant difference, as determined by one-way ANOVA with Tukey’s HSD tests (P□<□0.05). A quadratic function was fitted to the data as the approximation curve.

Under continuous low light (CLL), however, dry weight showed a convex relationship with PGR5 protein abundance but did not differ significantly among the lines and showed no clear relationship with PGR5 protein abundance (Fig. 4G).

Leaf area did not differ significantly among the lines under either FLL or CLL conditions (Figs. 4C, 4H).

In contrast, leaf mass per area (LMA) exhibited a clear relationship with PGR5 protein abundance under FLL conditions (Fig. 4D) indicating that the differences in plant biomass observed under FLL conditions were not associated with changes in leaf expansion. LMA showed a convex relationship with PGR5 protein level, with the highest value observed in the *PGR5ox 65* line. Under CLL conditions, LMA did not differ significantly among the lines (Fig. 4I).

Photosynthetic parameters measured after the growth treatments also showed line-dependent differences. Under FLL conditions, the *pgr5-5* mutants exhibited the lowest Fv/Fm values among the lines (Fig. 4E), indicating increased photoinhibition in the absence of PGR5. In contrast, under CLL conditions, the two highest-expressing lines (*PGR5ox 5-10* and *PGR5ox 25-10*) showed significantly lower Fv/Fm values than the other lines (Fig. 4J).

Pm showed a convex relationship with PGR5 protein abundance under both FLL and CLL conditions (Figs. 4F, 4K), with the highest values observed in the *PGR5ox 65* line. These results indicate that moderate accumulation of PGR5 enhances PSI stability, photosynthetic capacity and biomass production under fluctuating low-light environments.

#### 2) Growth responses under high-light conditions

Plants were also grown under HL conditions to evaluate line-dependent growth responses under stronger irradiance. After three weeks of treatment, the *pgr5-5* mutants showed significantly lower dry weight and leaf area than the other lines under both fluctuating high light (FHL) and continuous high light (CHL) conditions (Figs. 5B–5C; 5G–5H).

**Fig. 5.**
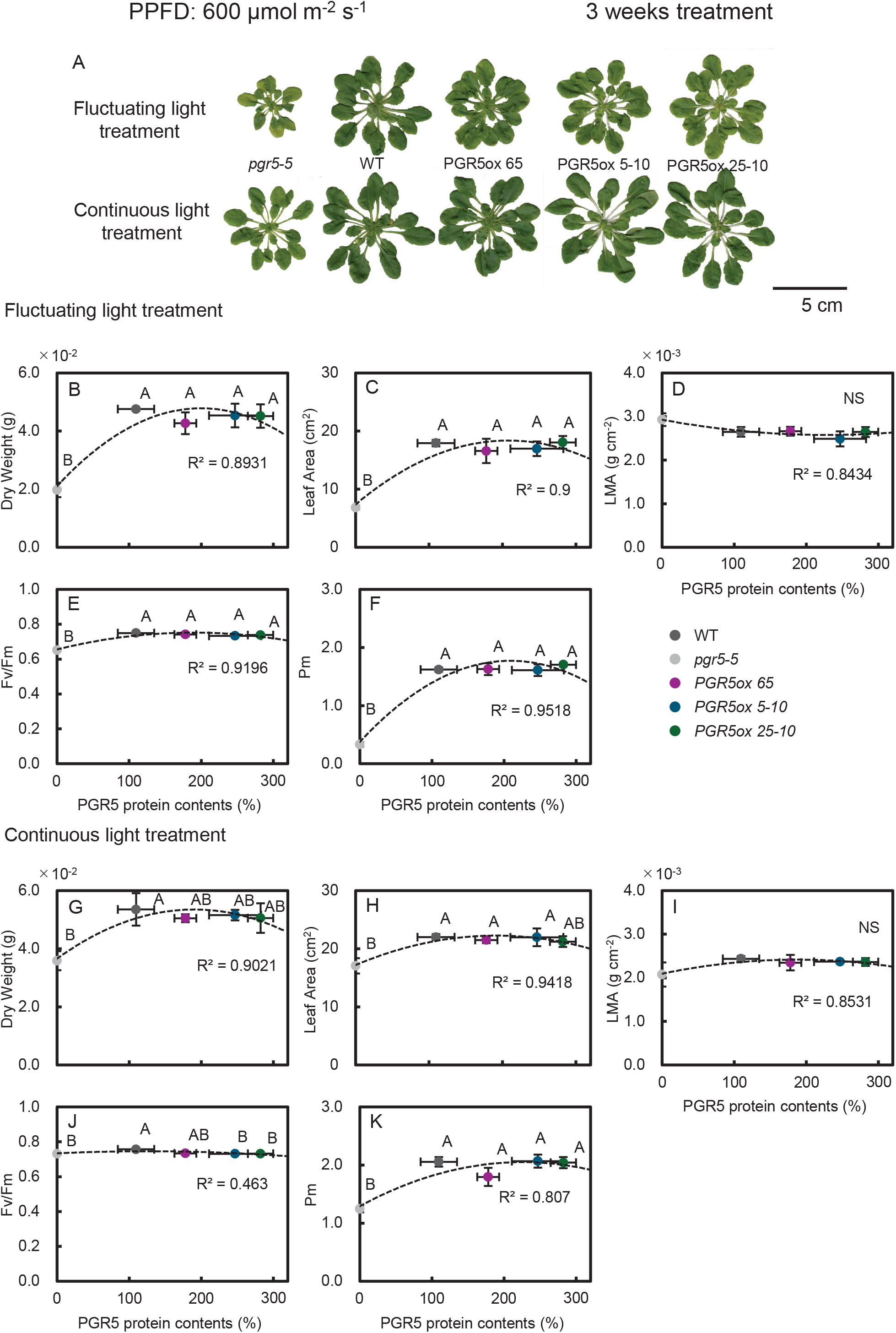
Plants Phenotype, growth analysis, chlorophyll fluorescence and P700 redox state under 3 weeks of FHL treatment in WT, *pgr5-5, PGR5ox 65, PGR5ox 5-10* and *PGR5ox 25-10*. **A**: Plants were cut and scanned after 3 weeks FHL treatment. Upper plants were grown under FHL condition and lower plants were grown under CHL condition. A plot of (**B**) Dry Weight, (**C**) Leaf Area, (**D**) LMA and (**E**) Fv/Fm as a function of PGR5 protein contents under FHL condition. A plot of(**F**) Dry Weight, (**G**) Leaf Area, (**H**) LMA and (**I**) Fv/Fm as a function of PGR5 protein contents under CHL condition. Data are means ± SE (n□=□4). For each point, different letters indicate a significant difference, as determined by one-way ANOVA with Tukey’s HSD tests (P□<□0.05). A quadratic function was fitted to the data as the approximation curve.

In contrast, leaf mass per area did not differ significantly among the lines under either FHL or CHL conditions (Figs. 5D, 5I), indicating that the reduced biomass observed in the *pgr5-5* mutants under HL conditions was mainly associated with reduced leaf development rather than changes in leaf structural traits.

Consistent with these growth differences, the physiological parameters Fv/Fm and Pm were significantly lower in the *pgr5-5* mutants than in the other lines under both FHL and CHL conditions (Figs. 5E–5F; 5J–5K), indicating reduced photosystem stability and PSI capacity in the absence of PGR5 under high-light environments.

### Time series analysis of ETRII and ETRI under photosynthetic induction

To further investigate induction kinetics, the first five minutes of the first and second induction phases were extracted from the time-course data (Fig. S5) and analyzed the time required for each parameter to reach 50% (t_50_), 60% (t_60_), 70% (t_70_), 80% (t_80_), and 90% (t_90_) of its maximum value after each step increase in light (Figs. S6, S7).

Under LL conditions, the measured ETRII values were similar among the lines during both induction phases (Figs. S5A–S5B). However, the *t*_*50*_–*t*_*90*_ values were larger in the highest PGR5-expressing lines (*PGR5ox 5-10* and *PGR5ox 25-10*), indicating slower PSII electron transport induction (Figs. S6A–S6B).

For PSI electron transport, the *pgr5-5* mutants showed lower ETRI values and slower induction kinetics compared with the other lines (Figs. S5C–S5D; Figs. S6C–S6D).

Under HL conditions, the *pgr5-5* mutants consistently exhibited the lowest ETRII, ETRI and *t*_*50*_–*t*_*90*_ values during the induction phases (Figs. S5E–S5H; S7A–S7D). These results indicate that PGR5 plays an essential role in maintaining efficient electron transport during rapid light transitions.

## Discussion

This study provides a comprehensive analysis of how varying levels of the PGR5 protein influence photosynthetic induction, PSI protection, and plant growth under fluctuating light conditions in *Arabidopsis thaliana*. Using a series of transgenic lines with graded PGR5 protein levels, we quantified chlorophyll fluorescence parameters, PSI redox dynamics, electron transport rates, and growth responses under both FLL and FHL regimes.

Our results revealed a non-linear relationship between PGR5 protein abundance and photosynthetic performance. Moderate PGR5 overexpression (approximately twofold, as in *PGR5ox 65*) enhanced photosynthetic induction (Fig. S4A) and resulted in increased biomass accumulation under FLL conditions (Figs. 4B–4D). In contrast, excessive overexpression (e.g., in *PGR5ox 5-10* and *PGR5ox 25-10*) resulted in impaired growth. Conversely, the *pgr5-5* mutants showed sluggish PSI electron transport (Figs. 2-1K, S4B), sustained PSI over-reduction (Fig. 2-1I), and reduced plant growth (Figs. 4B–4D, 4G–4I).

These findings highlight the central role of the PGR5 protein in dynamically adjusting photosynthetic electron flow through PSI during rapid light transitions. By facilitating CET, PGR5 contributes to maintaining PSI redox balance and protecting PSI from photodamage while sustaining carbon gain. Our results therefore extend previous studies demonstrating the importance of CET in photosynthetic regulation under fluctuating light conditions (Yamori and Shikanai, 2016).

### Differences in fluctuating light regimes explain the non-lethal phenotype of pgr5-5 in this study

Although the *pgr5-5* mutants exhibited clear physiological damage under fluctuating light conditions in this study, the phenotype was not lethal. Under long-term cultivation under FHL conditions, the *pgr5-5* mutants showed significantly lower dry weight, leaf area, Fv/Fm, and Pm compared with WT and the overexpression lines (Figs. 5B–5C; 5E–5F).

This outcome differs from previous studies reporting that *pgr5* mutants become lethal under fluctuating light (Tikkanen et al., 2010; Suorsa et al., 2012). We propose that this discrepancy largely reflects differences in the fluctuating light regimes employed. The term “fluctuating light” encompasses a wide range of light patterns, including stochastic light fluctuations that occur naturally in plant canopies as well as artificial square-wave regimes alternating between low and high light intensities.

In many previous studies reporting lethality in *pgr5*, the high-light phases were extremely intense and short in duration (e.g., one or two minutes), imposing severe physiological stress on the photosynthetic apparatus (Suorsa et al., 2012; Kono et al., 2014). In contrast, the fluctuating light regime used in the present study was relatively moderate. The high-light phase reached 600 µmol photons m□^2^ s□^1^, which is near the lower range of light saturation in *Arabidopsis thaliana*, and the high-light phase lasted 10 minutes.

These differences in light intensity and duration likely imposed milder PSI stress and therefore allowed the *pgr5* mutants to survive despite clear physiological impairment. Our results therefore highlight that the severity of the *pgr5* phenotype strongly depends on the characteristics of the fluctuating light environment.

### Optimized CET activity promotes photosynthetic induction and biomass accumulation

Our findings further demonstrate that optimized CET activity, achieved through moderate PGR5 protein accumulation, significantly improves photosynthetic induction and biomass production under fluctuating light conditions.

Lines with moderate PGR5 overexpression (e.g., *PGR5ox 65*) exhibited faster activation of both PSII and PSI electron transport, as indicated by shorter t□□values for ETRII and ETRI (Figs. S6, S7). These lines also displayed higher Y(I) and ETRI/ETRII ratios (Figs. 2-1G, 2-1L), suggesting enhanced PSI oxidation capacity and improved coordination between PSI and PSII electron transport.

Under FL conditions, PSI is particularly susceptible to over-reduction because downstream metabolic processes such as the Calvin–Benson cycle require time to become fully activated. Previous studies have shown that PGR5-dependent CET contributes to PSI protection through both donor-side regulation at the Cyt *b*□*f* complex (photosynthetic control) and acceptor-side regulation of PSI redox poise (Suorsa et al., 2012; Yamamoto and Shikanai, 2019; Zhou et al., 2022; Kono et al., 2025).

Our results extend these findings by demonstrating that moderate enhancement of CET capacity not only alleviates PSI photoinhibition but also promotes downstream CO□assimilation and biomass accumulation.

Conversely, the *pgr5-5* mutants lacked CET activity and therefore exhibited sluggish PSI electron transport (Fig. S4B), reduced ETRI (Fig. 2-1K), and sustained PSI over-reduction (Y(NA); Fig. 2-1I). These results are consistent with previous studies demonstrating the critical role of CET in protecting PSI from photoinhibition (Munekage et al., 2002; Yamori and Shikanai, 2016).

### Excessive CET activity disturbs photosynthetic regulation

Interestingly, although moderate PGR5 overexpression improved photosynthetic performance, excessively high PGR5 expression did not confer additional benefits. Instead, the highest-expressing lines (*PGR5ox 5-10* and *PGR5ox 25-10*) showed reduced Fv/Fm (Fig. 1D), slower Y(II) induction (Fig. 2-1D), and excess NPQ induction (Fig. 2-1E).

These responses suggest that excessive CET activity may lead to excessive lumen acidification and premature activation of photoprotective energy dissipation mechanisms, thereby reducing photochemical efficiency. In addition, transient spikes of NPQ, Y(ND) and Y(NA) were observed in these lines under high-light conditions (Fig. S3A, S3C, S3D), indicating disturbances in redox homeostasis. These results are consistent with previous studies demonstrating that the activation of the Calvin cycle is delayed in PGR5-overexpressing lines (Okegawa et al., 2022).

These results indicate that CET activity must be finely balanced to optimize photosynthetic regulation. Both insufficient CET (as in *pgr5-5* mutants) and excessive CET (as in high PGR5 overexpression lines) lead to suboptimal physiological outcomes.

### Implications for optimizing photosynthesis in fluctuating light environments

Collectively, our results demonstrate that precise tuning of PGR5 protein abundance is essential for maintaining optimal photosynthetic performance. In wild-type plants, this balance is naturally maintained by PGRL1 and PGRL2, which prevent the excessive accumulation or degradation of PGR5 (Rühle et al., 2021). Moderate enhancement of CET capacity improved photosynthetic induction, stabilized PSI redox balance, and promoted biomass accumulation under FLL environments.

Because natural light environments within plant canopies are highly dynamic, particularly under shaded conditions, optimizing CET activity may represent a promising strategy for improving crop productivity in field environments. Our findings therefore provide new insights into the physiological significance of CET regulation and highlight the importance of balancing photoprotection and energy conversion efficiency in dynamic light environments.

## Conclusion

This study demonstrates that the level of PGR5 protein is a key determinant of photosynthetic induction efficiency, PSI stability, and plant growth under fluctuating light environments. Moderate overexpression of PGR5 enhanced photosynthetic induction and biomass accumulation under fluctuating low light, whereas both insufficient and excessive PGR5 levels disrupted photosynthetic regulation.

These findings reveal that an optimal range of CET capacity is required to maximize photosynthetic performance and plant fitness under dynamic light conditions. Fine-tuning PGR5-dependent CET may therefore represent a promising strategy for improving photosynthetic efficiency and crop productivity in naturally fluctuating light environments.

## Materials & methods

### Plant materials and cultivation

We grew *Arabidopsis thaliana* wild type (ecotype Columbia *gl1*, WT), *PGR5* knockout mutant (*pgr5-5)*, and four transgenic lines overexpressing *PGR5* under the control of the cauliflower mosaic virus 35S promoter (*35S::PGR5*). The line names are *PGR5ox 20, PGR5ox 65, PGR5ox 5-10*, and *PGR5ox 25-10*, and they have different PGR5 protein levels. The knockout mutant was generated by Kobayashi et al. (2024), and these overexpression lines were generated by Okegawa et al. (2007).

All plants were grown in 400 ml pots or cell trays containing a 1:1 mixture of soil (Metro-Mix 350J; Hyponex Japan, Osaka, JAPAN) and expanded vermiculite. They were cultivated under growth chamber conditions (120 μmol photons m^−2^ s^−1^, 10 h light/14 h dark cycles, 60 %RH and 23 °C). The plants in pots were cultivated for 6 weeks, and Hyponex liquid fertilizer (Hyponex Japan, OSAKA, JAPAN) was applied in week 4. These plants were then subjected to the photosynthetic induction measurements described below. The *pgr5-5* mutant was cultivated and measured at a different time from other lines.

Five lines of plants (WT, *pgr5-5, PGR5ox 65, PGR5ox 5-10* and *PGR5ox 25-10*) in cell trays were cultivated in continuous light for 3 weeks and then transferred to fluctuating light cultivation systems. We installed two systems in a room where the temperature was kept at 23 °C, and one was set to an FL condition (10 min. high light/10 min. low light cycles during the light period) and the other to continuous light (CL) condition. The total photon flux was kept the same between the two light conditions. The light cycles for these conditions were 10 h light/14 h dark. Two light intensities (600 μmol photons m^−2^ s^−1^ and 100 μmol photons m^−2^ s^−1^) were available as high-light, and the low-light was set to 10 μmol photons m^−2^ s^−1^. In summary, the cultivation system included four light conditions: FLL (100 and 10 μmol photons m^−2^ s^−1^), CLL (average intensity of FLL), FHL (600 and 10 μmol photons m^−2^ s^−1^), CHL (average intensity of FHL). We placed two cell trays in each system for 3 weeks, after which they were subjected to the chlorophyll fluorescence and P700 redox measurements described below.

### Measurement of chlorophyll fluorescence and P700 in a photosynthetic induction response

Chlorophyll fluorescence and P700 were measured simultaneously with a Dual-PAM-100 measuring system (Walz, Effeltrich, Germany), as described previously (Qu et al., 2025; Kono et al., 2025). Measurements were performed on fully expanded, uppermost leaves of 6-week-old plants. Leaves were dark-adapted for a day, and then a saturating pulse was applied to obtain F_v_/F_m_ (Genty et al., 1989) and Pm (Klughammer and Schreiber, 1994; Klughammer and Schreiber, 2008b). Chlorophyll fluorescence was determined with a Dual-PAM-100’s chamber. First, a saturating pulse was applied to plant leaves to obtain maximum fluorescence. After measuring of the quantum yield of PSII (Y(II)), we calculated the electron transport rate through PSII (ETRII) as:

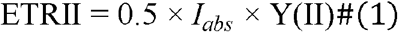

The P700 was also determined with the Dual-PAM-100’s chamber. First, leaves were treated with a saturating pulse and far-red light to obtain P_m_. After measurement of the quantum yield of PSI (Y(I)), we calculated the electron transport rate through PS I (ETRI) as:

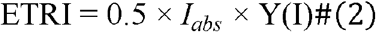

where 0.5 is the fraction of absorbed light allocated to each photosystem, and *I*_*abs*_ is the absorbed irradiance, taken as 0.84 of the incident irradiance. In addition, NPQ (Bilger and Björkman, 1990), 1−qL (Kramer et al., 2004), Y(ND) (Klughammer and Schreiber, 2008b), and Y(NA) (Klughammer and Schreiber, 2008b) were calculated. NPQ represents the extent of non-photochemical quenching, whereas 1−qL reflects the redox state of the plastoquinone pool, specifically the primary electron acceptor Q_A_. Y(ND) is the quantum yield of non-photochemical energy dissipation due to donor side limitation, and Y(NA) is the quantum yield of non-photochemical energy dissipation due to acceptor-side limitation.

Chlorophyll fluorescence and P700 were measured every 60 seconds at a CO_2_ concentration of 400 μmol mol^−1^, 60 %RH, and 23.0 °C. Two distinct light conditions were employed during the measurements: a HL condition and a LL condition. In the HL condition, the actinic light was initially turned off for 3 minutes, then increased sequentially to 10 μmol photons m^−2^ s^−1^ for 5 minutes and 600 μmol photons m^−2^ s^−1^ for 30 minutes. It was then reduced to 10 μmol photons m^−2^ s^−1^ for 5 minutes, raised again to 600 μmol photons m^−2^ s^−1^ for 10 minutes, and finally returned to 0 μmol photons m^−2^ s^−1^ for 5 minutes. In contrast, under the LL condition, the light intensity followed the same temporal pattern but with a lower peak irradiance of 100 μmol photons m^−2^ s^−1^.

For the investigation of photosynthetic induction, plants were selected at random and measured from 08:00 to 16:00 to avoid confounding samples with the time of day and to minimize any diurnal influences. The day before measurements, plants were removed from the growth chamber before the light period began and were kept in a dark room at an air temperature of 25 °C, relative humidity of 65% and ambient CO_2_ concentration. A leaf was then placed into the measuring chamber whilst maintaining dark conditions, with an air temperature of 23 °C, relative humidity of 60% and 400 µmol CO_2_ mol^−1^.

We evaluated t_50_–t_90_ of ETRII and ETRI after each step increase in light. Before evaluating t_50_–t_90_, the plot sequences of ETRII and ETRI were fitted to a quadratic function. After confirming these curve fittings, we calculated t_50_–t_90_ using these regression curves to determine their accurate values.

### Measurement of chlorophyll fluorescence, P700, SPAD value and growth analysis under a long-term fluctuating light condition

Chlorophyll fluorescence and P_700_ were also measured simultaneously with a Dual-PAM-100 measuring system. The measurements were conducted immediately following harvest. All measurements were performed under controlled environmental conditions maintained at 25 °C, 65 %RH, and ambient CO□concentration. First, F_v_/F_m_ was determined, followed by the measurement of P_m_.

Following these measurements, the plants were subjected to growth analysis. The entire shoot of each plant was first scanned at 600 dpi using a flatbed scanner (CanoScan LiDE 220, Canon Inc., Tokyo, Japan) to create a digital image. The total leaf area was subsequently calculated from these images using ImageJ software (National Institutes of Health, MD, USA). After scanning, each plant was placed into an individual paper envelope and dried in an oven at 80°C for three days. The total dry mass was then determined by weighing each dried sample. Finally, the Leaf Mass per Area (LMA) was calculated by dividing the total dry mass by the leaf area.

### Statistical analysis

Differences among three or more groups were analysed by ANOVA using the Tukey–Kramer test according to Sakoda et al. (2021). Pearson’s correlation coefficient (r) was calculated, and the significance of relationships was tested by two-sided t-tests (P < 0.05). All statistical analyses were conducted in Python 3.13.4 (Python Software Foundation, DE, USA).

### Western-blotting of PGR5 and PetC proteins and SDS-PAGE of RubisCO

Leaf tissues were sampled after chlorophyll fluorescence and P700 measurements, frozen in liquid nitrogen and stored at −80 °C until analysis. The RubisCO content was determined by formamide extraction of SDS-PAGE separated, Coomassie Brilliant Blue R-250 stained bands corresponding to the large and small subunits of RubisCO as described in Qu et al. (2021). The proteins were separated by 16.5 % SDS-PAGE using the conventional Tris-tricine buffer system for PGR5 and PetC proteins (Schägger and von Jagow, 1987) and transferred onto a PVDF (polyvinylidene fluoride) membrane. Immunodetection with antibodies against PGR5 and PetC was performed as described in Munekage et al. (2002) and Okegawa et al. (2007). For immunoblot analysis, extracts from four WT individuals were pooled based on their chlorophyll content to create a ‘WT mix’, which was used as a protein loading control.

## Data Availability Statement

The data supporting this work are available within the paper and its Supporting Information.

## Acknowledgments and Funding

This work was supported by JSPS KAKENHI (Grant Number 21H02171 to W.Y.), JST ALCA-Next (Grant Number JPMJAN25D2 to W.Y.), and the Sasagawa Scientific Research Grant from The Japan Science Society (Grant Number 2024-4013 to K.T.).

## Author Contributions

K.T. and W.Y. conceived and designed the study. K.T., H.K., Y.O. and W.Y. conducted experiments and performed data analysis. K.T., Y.O., T.S. and W.Y. wrote the manuscript. All authors discussed the results and revised the manuscript.

## Figure Legend

***Fig. S1.***

Fig. S1. Protein accumulation in WT, *pgr5-5, PGR5ox 20, PGR5ox 65, PGR5ox 5-10* and *PGR5ox 25-10*. **A**: Relative amount of PetC protein upon WT being set at 100%. **B**: Relative amount of RubisCO Large subunit protein upon WT being set at 100%. Data are means ± SE (n□≥□3), and dots indicate individual data points. For each point, different letters indicate a significant difference, as determined by one-way ANOVA with Tukey’s HSD tests (P□<□0.05).

***Fig. S2.***

Fig. S2. Chlorophyll fluorescence and P700 analysis of a photosynthetic induction response in WT, *pgr5-5, PGR5ox 20, PGR5ox 65, PGR5ox 5-10* and *PGR5ox 25-10* under the LL condition. Time series changes in (**A**) NPQ, (**B**) 1−qL, (**C**) Y(ND), Y(NA), (**E**) ETRII, (**F**) ETRI and (**G**) ETRI/ETRII ratio. Data are means ± SE (n□≥□4)

***Fig. S3.***

Fig. S3. Chlorophyll fluorescence and P700 analysis of a photosynthetic induction response in WT, *pgr5-5, PGR5ox 20, PGR5ox 65, PGR5ox 5-10* and *PGR5ox 25-10* under the HL condition. Time series changes in (**A**) NPQ, (**B**) 1−qL, (**C**) Y(ND), (**D**) Y(NA), (**E**) ETRII, (**F**) ETRI and (**G**) ETRI/ETRII ratio. Data are means ± SE (n□≥□4)

***Fig. S4.***

Fig. S4. Chlorophyll fluorescence and P700 analysis of a photosynthetic induction response in WT, *pgr5-5, PGR5ox 20, PGR5ox 65, PGR5ox 5-10* and *PGR5ox 25-10*. A plot of (**A**) t_50_ of ETRII under the LL condition, (**B**) t_50_ of ETRI under the LL condition, (**C**) t_50_ of ETRII under the HL condition, and (**D**) t_50_ of ETRI under the HL condition as a function of PGR5 protein contents in the 1^st^ induction. Data are means ± SE (n□≥□4). For each point, different letters indicate a significant difference, as determined by one-way ANOVA with Tukey’s HSD tests (P□<□0.05). The regression line was obtained using the method of least squares and evaluated by calculating the R^2^ and p-value based on Pearson’s correlation coefficient.

***Fig. S5.***

Fig. S5. Electron transport rate during the first 5 minutes of the 1^st^ and 2^nd^ induction phases in WT, *pgr5-5, PGR5ox 20, PGR5ox 65, PGR5ox 5-10* and *PGR5ox 25-10* under both the LL and HL condition. Time series changes in (**A**) ETRII of 1^st^ induction under LL, (**B**) ETRII of 2^nd^ induction under LL, (**C**) ETRI of 1^st^ induction under LL, (**D**) ETRI of 2^nd^ induction under LL, (**E**) ETRII of 1^st^ induction under HL, (**F**) ETRII of 2^nd^ induction under HL, (**G**) ETRI of 1^st^ induction under HL and (**H**) ETRI of 2^nd^ induction under HL. A quartic function was fitted to the data as the approximation curve.

***Fig. S6.***

Fig. S6. t_50_, t_60_, t_70_, t_80_ and t_90_ of (**A**) ETRII of 1^st^ induction, (**B**) ETRII of 2^nd^ induction, (**C**) ETRI of 1^st^ induction and (**D**) ETRI of 2^nd^ induction under the LL condition. Data are means ± SE (n□≥□4). For each point, different letters indicate a significant difference, as determined by one-way ANOVA with Tukey’s HSD tests (P□<□0.05).

***Fig. S7.***

Fig. S7. t_50_, t_60_, t_70_, t_80_ and t_90_ of (**A**) ETRII of 1^st^ induction, (**B**) ETRII of 2^nd^ induction, (**C**) ETRI of 1^st^ induction and (**D**) ETRI of 2^nd^ induction. under the HL condition. Data are means ± SE (n□≥□4). For each point, different letters indicate a significant difference, as determined by one-way ANOVA with Tukey’s HSD tests (P□<□0.05).

