## Supplementary figures and images for "Moderate overexpression of *PROTON GRADIENT REGULATION 5* improves photosynthetic performance and plant growth under fluctuating low light in *Arabidopsis thaliana*"

### Supplemental Figure 1

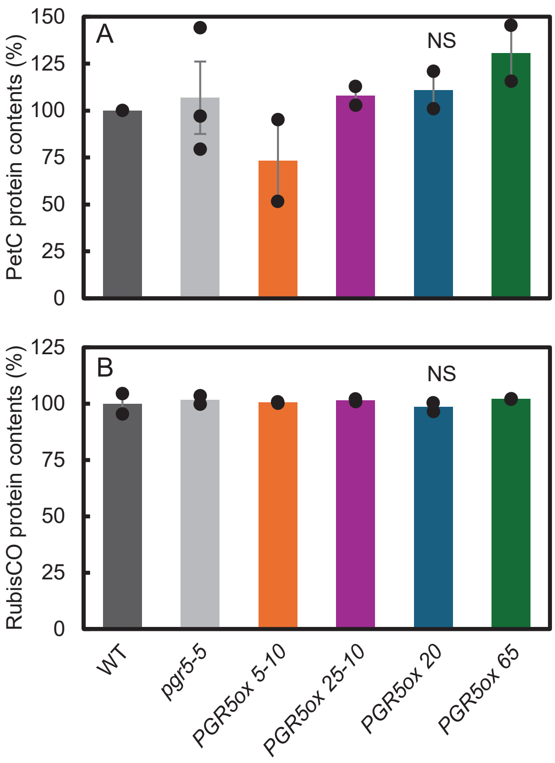

### Supplemental Figure 2

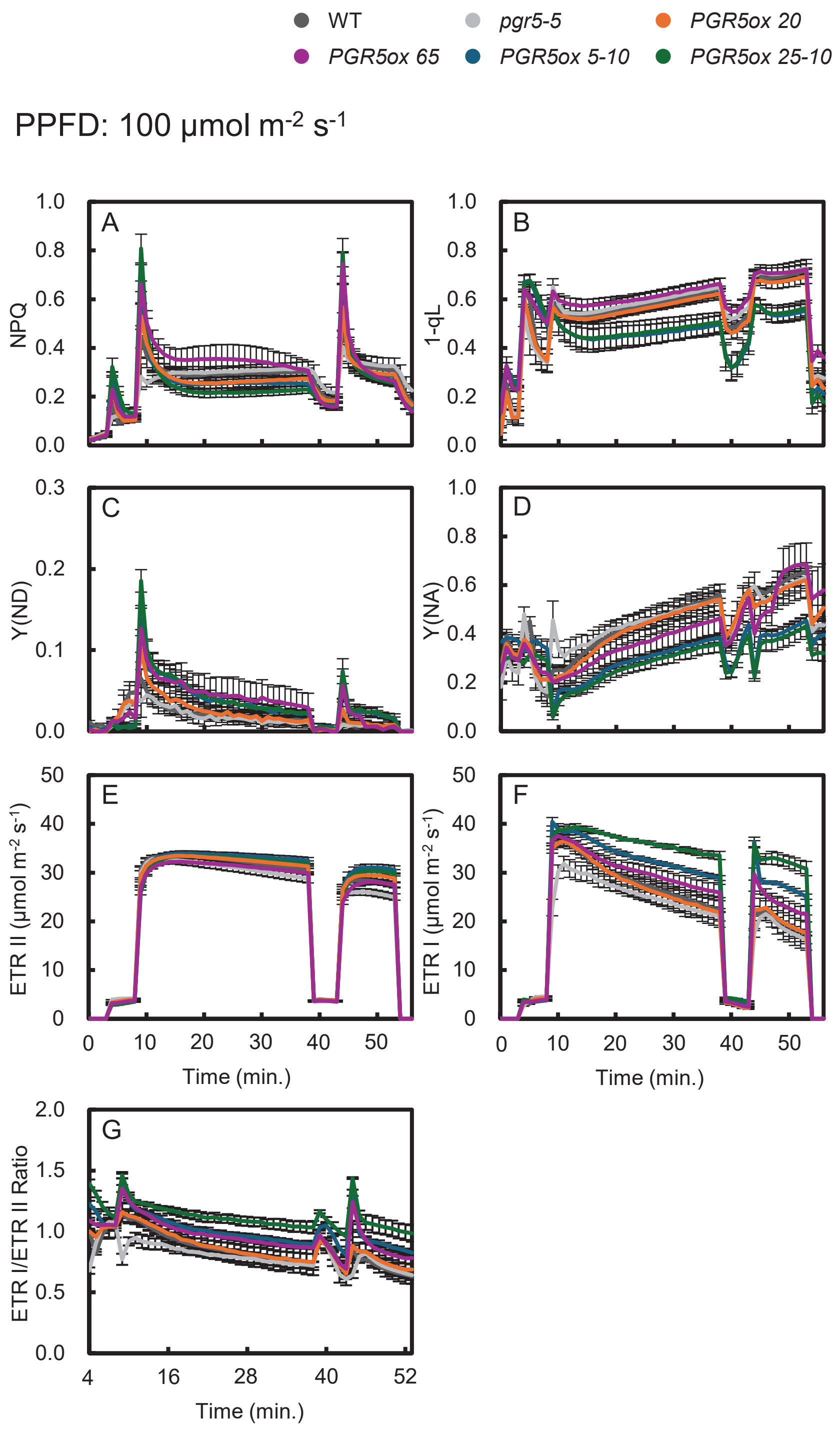

### Supplemental Figure 3

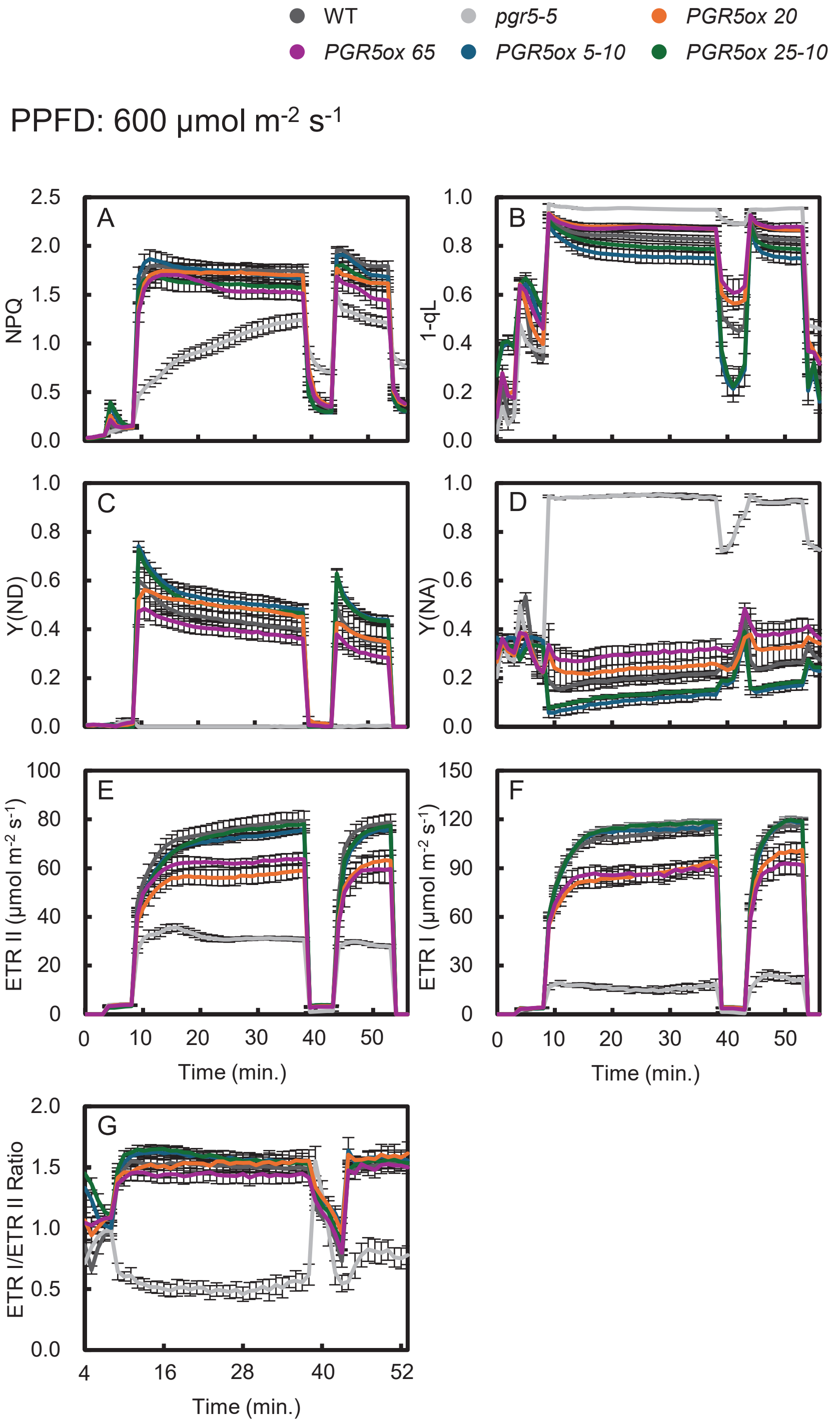

### Supplemental Figure 4

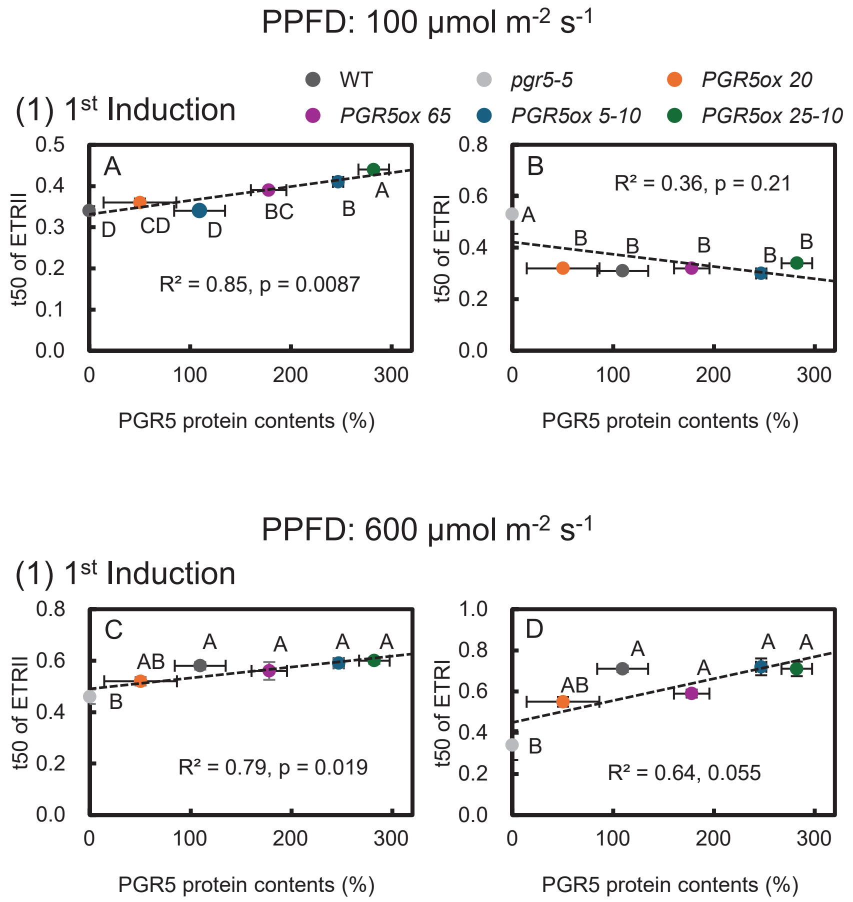

### Supplemental Figure 5

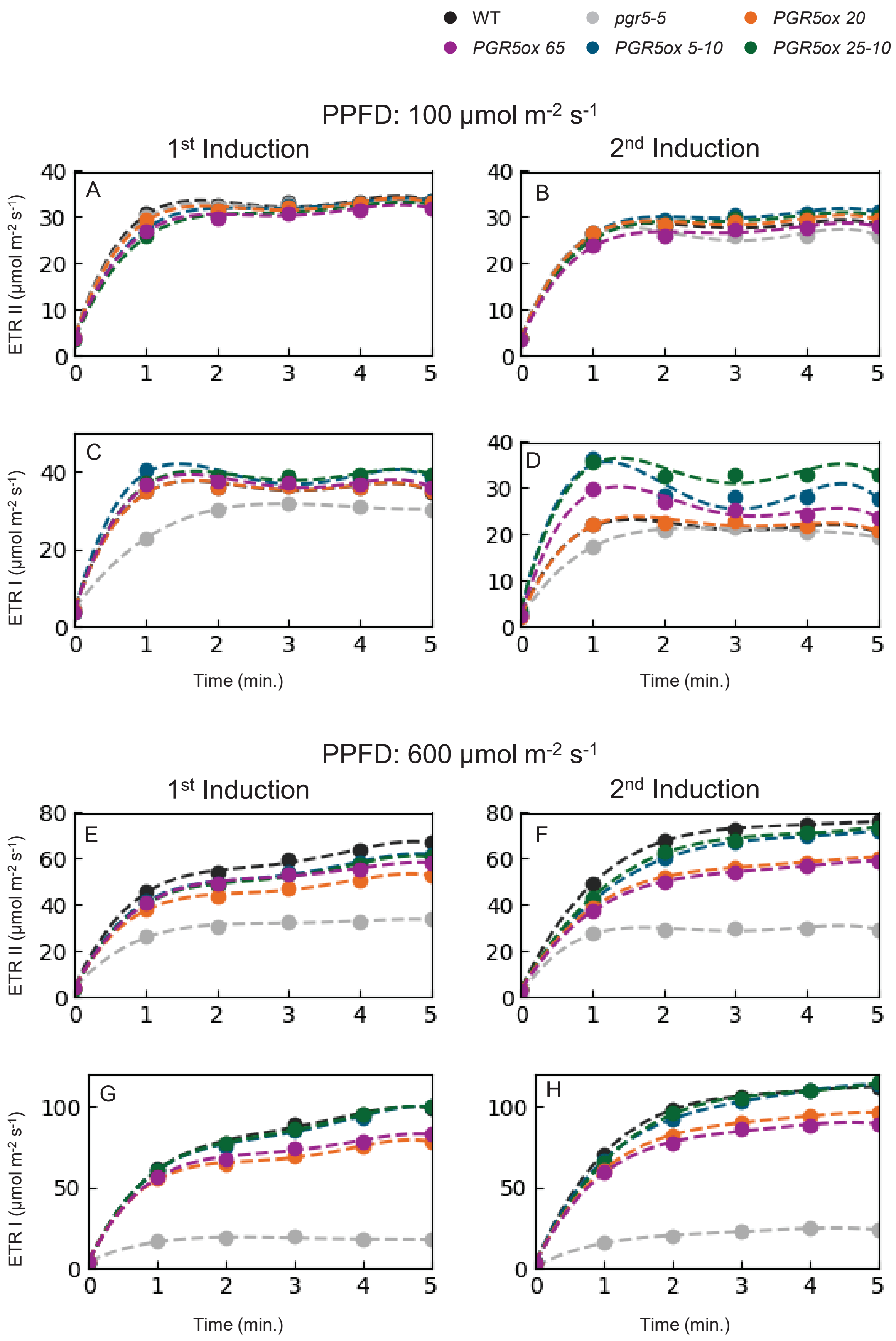

### Supplemental Figure 6

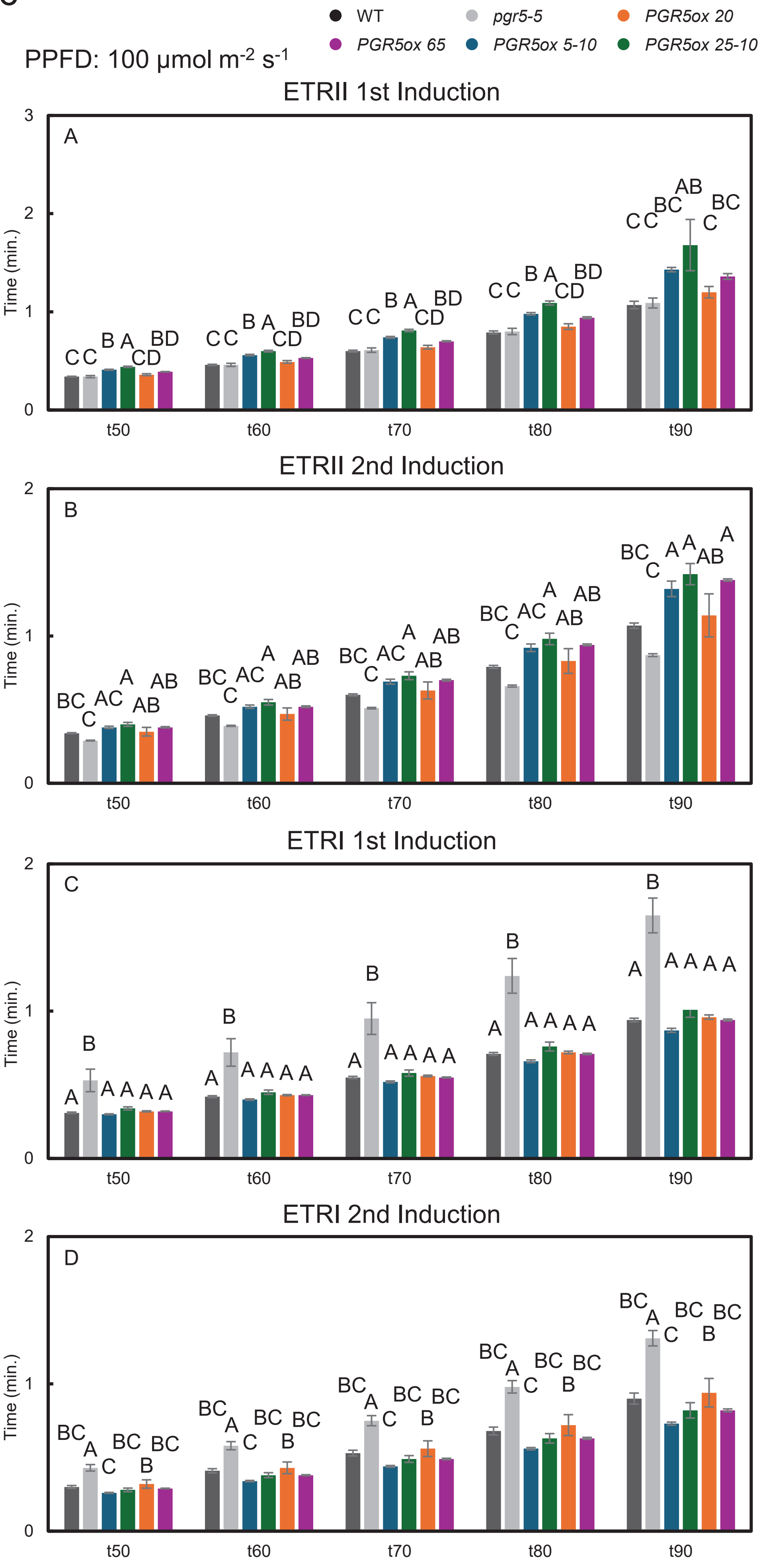

### Supplemental Figure 7

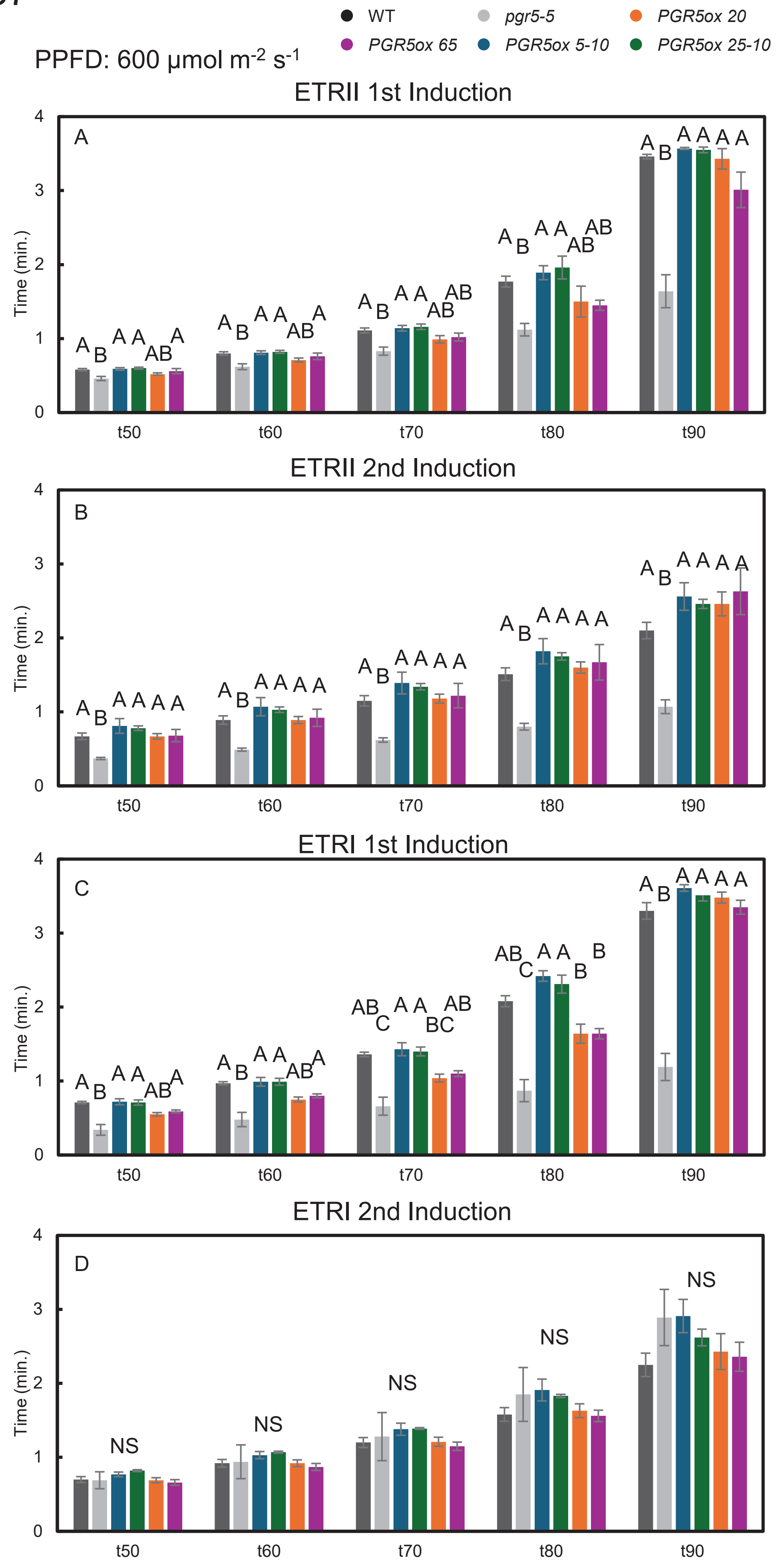
